# The neural basis of resting-state fMRI functional connectivity in fronto-limbic circuits revealed by chemogenetic manipulation

**DOI:** 10.1101/2023.06.21.545778

**Authors:** Catherine Elorette, Atsushi Fujimoto, Frederic M. Stoll, Satoka H. Fujimoto, Lazar Fleysher, Niranjana Bienkowska, Brian E. Russ, Peter H. Rudebeck

**Author notes:** These authors contributed equally to this work (Catherine Elorette, Atsushi Fujimoto). Joint last author (Brian E. Russ, Peter H. Rudebeck).

## Abstract

Measures of fMRI resting-state functional connectivity (rs-FC) are an essential tool for basic and clinical investigations of fronto-limbic circuits. Understanding the relationship between rs-FC and neural activity in these circuits is therefore vital. Here we introduced inhibitory designer receptors exclusively activated by designer drugs (DREADDs) into the macaque amygdala and activated them with a highly selective and potent DREADD agonist, deschloroclozapine. We evaluated the causal effect of activating the DREADD receptors on rs-FC and neural activity within circuits connecting amygdala and frontal cortex. Interestingly, activating the inhibitory DREADD increased rs-FC between amygdala and ventrolateral prefrontal cortex. Neurophysiological recordings revealed that the DREADD-induced increase in fMRI rs-FC was associated with increased local field potential coherency in the alpha band (6.5-14.5Hz) between amygdala and ventrolateral prefrontal cortex. Thus, our multi-disciplinary approach reveals the specific signature of neuronal activity that underlies rs-FC in fronto-limbic circuits.

## INTRODUCTION

Resting-state functional connectivity (rs-FC) is defined as the temporal correlation of activity signals between two brain regions in the absence of a stimulus or task (Chen et al., 2020). In the three decades since the invention of functional magnetic resonance imaging (fMRI), and the development of resting-state analyses, rs-FC has become a standard tool for examining network-level functional variations in the brain. In particular, it has been used to track variation in frontal circuits in development (Grayson and Fair, 2017), aging (Rosenberg et al., 2020), as well as neurological and psychiatric disease (Greicius, 2008). Indeed, in psychiatric research, patterns of rs-FC in fronto-limbic circuits have become increasingly used as biomarkers of different subtypes of depression (Chai et al., 2023), autism (Ren et al., 2023), anxiety (Li et al., 2023), and schizophrenia (Zhu et al., 2016). The validation and use of these biomarkers hold the potential to enable personalized treatments for subtypes of psychiatric disorders (Yamada et al., 2017; Lee et al., 2022).

Despite the widespread use of rs-FC in basic and clinical research, the neural basis of these measures in fronto-limbic circuits is not clear. In sensory cortex, measures of fMRI rs-FC correlate with the temporal similarity or coherence of neural activity between distributed brain areas (He et al., 2008; Shmuel and Leopold, 2008; Schölvinck et al., 2010; Pan et al., 2013; Wilson et al., 2016). Specifically, several studies have observed a close relationship between fMRI rs-FC and local field potential (LFP) coherence in low-frequency bands (Leopold et al., 2003; He et al., 2008; Wang et al., 2012; Wilson et al., 2016; Wu et al., 2017). These findings have led to the widely accepted notion that this neural mechanism underlies rs-FC across the whole brain, including in fronto-limbic circuits. Recent studies of fMRI resting-state networks as well as neurophysiology indicate, however, that frontal areas exhibit functional timescales that are distinct from those in sensory areas (Murray et al., 2014; Manea et al., 2022). This potentially calls into question whether neural markers of rs-FC observed in sensory areas can be extrapolated to interactions between frontal cortex and limbic areas. In addition, many of the prior investigations of the basis of rs-FC focused on correlating fMRI signals with neural activity markers of functional communication, but did not assess how perturbing neural activity altered this relationship (Leopold et al., 2003; Shmuel and Leopold, 2008; Pan et al., 2011, 2013; Hutchison et al., 2015). Consequently, it is not known if manipulating neural activity in one part of the limbic system is associated with changes in rs-FC *and* corresponding changes in LFP coherency.

Here we set out to determine the basis of rs-FC in fronto-limbic circuits using a multi-disciplinary approach wherein we causally manipulated neural activity using Designer Receptors Exclusively Activated by Designer Drugs (DREADDs, **Figure 1**; Armbruster et al., 2007). Such chemogenetic approaches enable the reversible control of specific neuronal populations, can be combined with fMRI as well as neurophysiology, and are thus highly amenable for studying the basis of rs-FC. As a relatively new technology, there is conflicting evidence about the overall effect of DREADD-mediated inactivation at the systems level. A previous study in macaques found network-level decreases in rs-FC after the inactivation of the amygdala using clozapine-N-oxide (CNO; Grayson et al., 2016). However more recent rodent research has found a paradoxical increase in rs-FC following inactivation of the prefrontal cortex (Rocchi et al., 2022). Through the use of more refined techniques, we aim to reconcile these differences in the macaque.

**Figure 1.**
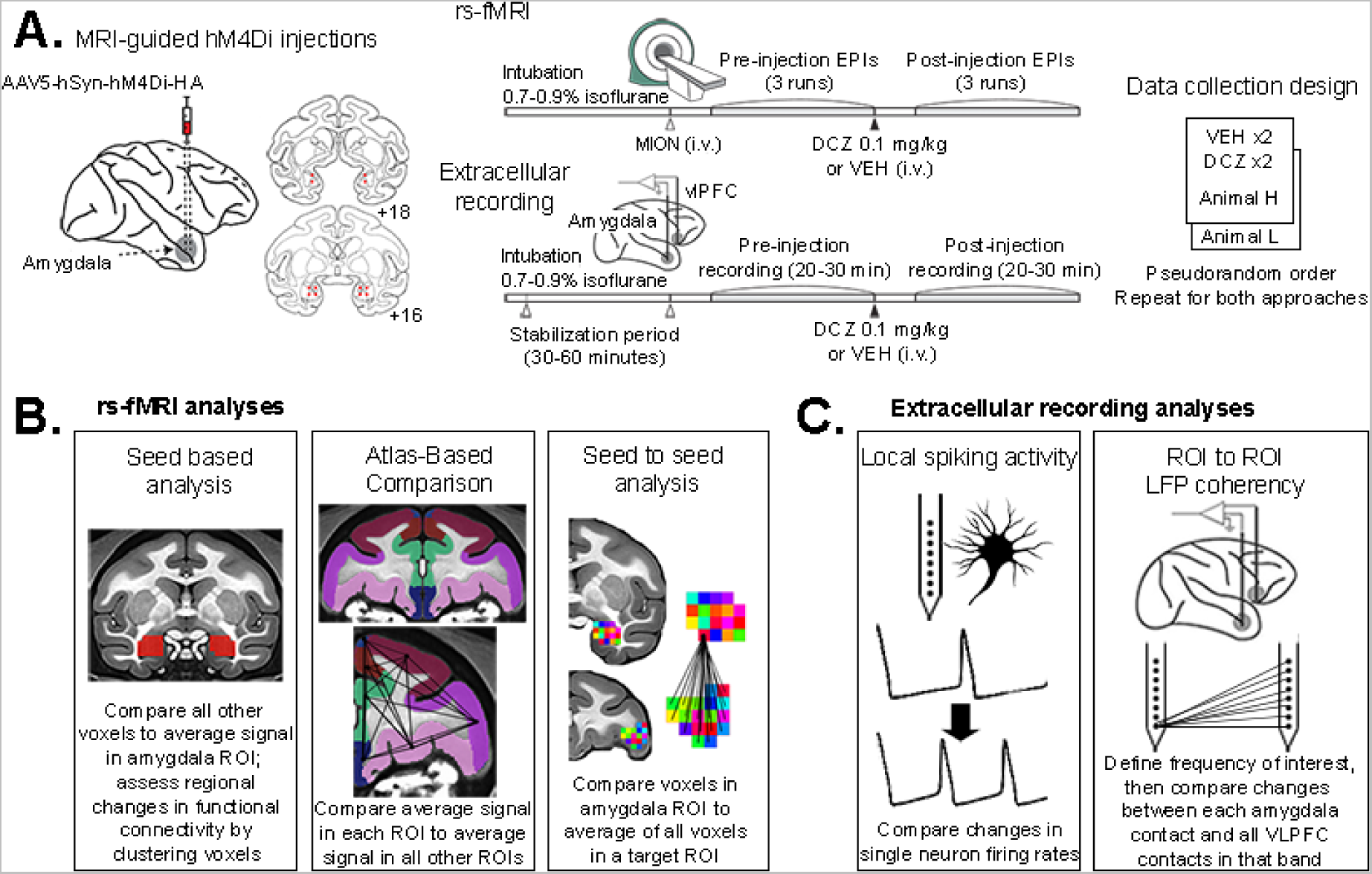
Schematic of experimental design and analysis approach. **A)** Experimental timeline from injection of DREADDs into amygdala, through data collection of fMRI and extracellular recordings. Types of **B)** fMRI or **C)** neural activity analyses performed.

Because of its central role in a host of psychiatric disorders, we specifically targeted inhibitory DREADDs (hM4Di) to the basolateral nucleus of the amygdala. We then assessed how manipulating activity in this area altered rs-FC and LFP coherency in fronto-limbic circuits. To aid selective manipulation of DREADD-expressing cells and assess the causal influence of their projections across the brain, we used deschloroclozapine (DCZ). This actuating drug is proven to be inert and has no effect on whole-brain functional connectivity (Fujimoto et al., 2022; Cushnie et al., 2023) or on various types of behaviors (Nagai et al., 2020; Upright and Baxter, 2020; Mimura et al., 2021). DREADD-mediated inhibition of the amygdala increased the brain-wide rs-FC of this area, and specifically increased rs-FC between amygdala and frontal cortex. Additionally, this increase in amygdala-frontal rs-FC was associated with increased LFP coherence, in specific frequency bands. Thus, our data provide direct evidence for a specific neural mechanism that underlies rs-FC in the circuits that connect the limbic system and frontal cortex.

## RESULTS

### Viral transfection of hM4Di DREADDs produced extensive coverage of amygdala nuclei

To examine how measures of fMRI rs-FC and neural activity within fronto-limbic circuitry relate to one another after focal inhibition of amygdala activity, we used a chemogenetic approach (**Figure 1A**). Two male macaques underwent a neurosurgery to transfect DREADD receptors (Armbruster et al., 2007) into the amygdala. We injected an AAV vector encoding an inhibitory DREADD (hM4Di) for pan-neuronal expression with a hemagglutinin (HA) marker protein. Virus injections were made bilaterally at 6 sites (3ul each) and were targeted towards the basolateral division of the amygdala (Amaral et al., 1992; Chareyron et al., 2011). Immunohistochemical analysis targeted to the HA marker protein revealed robust expression throughout the amygdala (**Figure 2A**). In both animals, the strongest labeling occurred in basal and basal accessory nuclei. Both animals additionally displayed labeling of cell bodies and axon terminals in the lateral nucleus, which varied by animal and by hemisphere. Specifically, animal H displayed a pattern of labeling that was concentrated medially and anteriorly within the amygdala. The periamygdaloid complex was well labeled in both hemispheres. Some sparse labeling of cell bodies was observed in central nucleus in the right hemisphere. In animal L, labeling was more medial in the left hemisphere and more lateral in the right hemisphere. In this animal, the left periamygdaloid complex showed strong labeling, while the right lateral and central nuclei were also well labeled. Further, labeled axon terminals were also visible in both animals throughout the amygdala, as well as in regions known to receive projections from basal amygdala, such as the ventral frontal cortex (**Figure 2C**). Overall, DREADD receptors were expressed bilaterally in the amygdala in both monkeys, with the greatest overlap between subjects in our target, the basal nuclei, which contains cortical projection neurons (Aggleton and Mishkin, 1984; Carmichael and Price, 1995).

**Figure 2.**
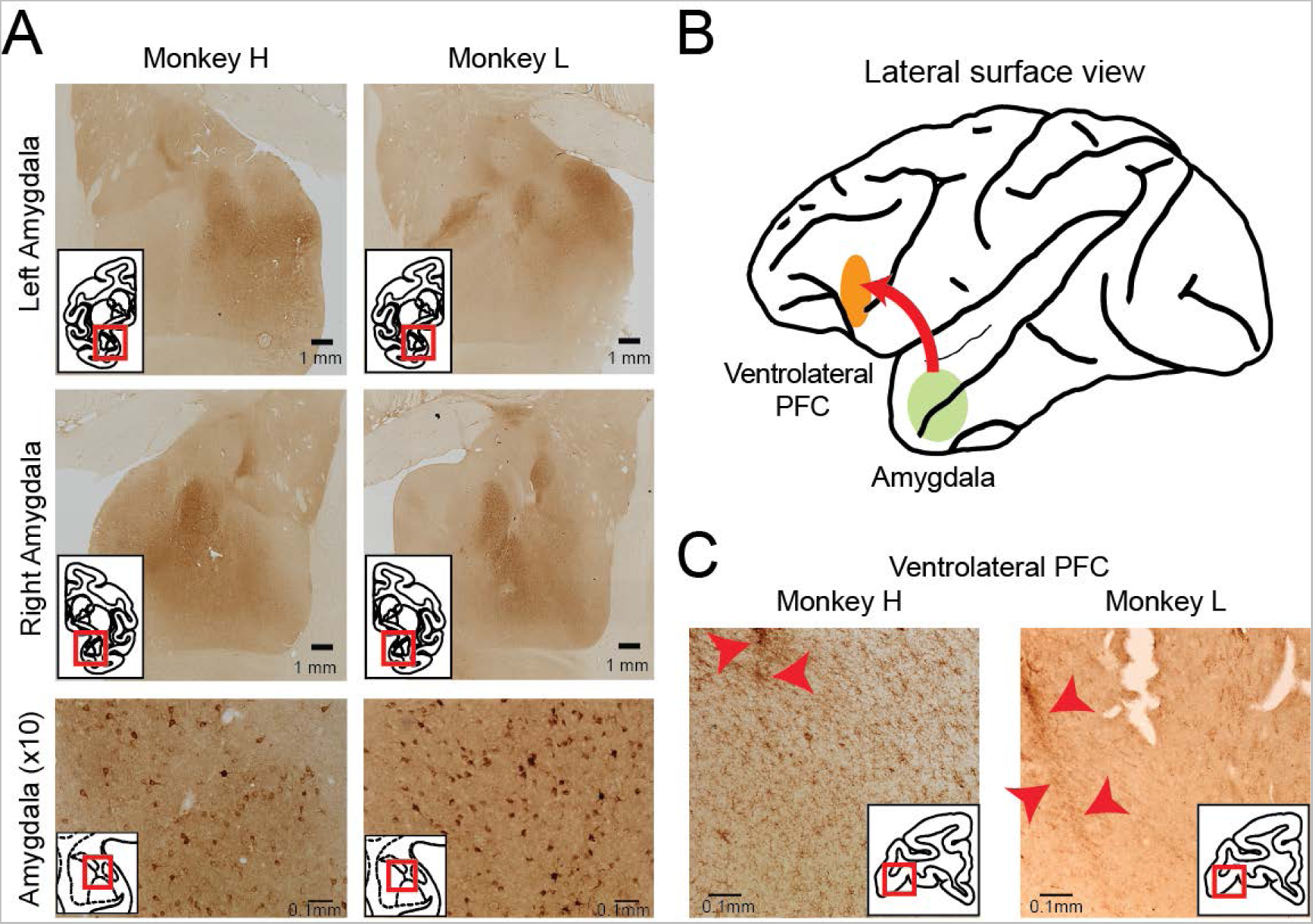
DREADD transfection confirmed by histological processing. **A)** Brain sections processed for HA tag showing representative images shown of whole amygdala (tiled images, top and middle) as well as DREADD transfected amygdala neurons (bottom). **B)** Schematic of DREADD injection site in amygdala and labeling of axonal projections in vlPFC. **C)** Brain sections processed for HA tag, showing labeling of axon terminals in vlPFC (see inset). Representative images are shown for each animal of the hemisphere in which extracellular recording of vlPFC occurred. Red arrows indicate electrode tracks.

### Increased amygdala rs-FC after chemogenetic inhibition

To determine the effects of chemogenetic inhibition of the amygdala on resting state networks in macaques, both animals underwent functional neuroimaging after amygdala DREADDs transfection. Following our previous imaging protocol, we maintained animals on light isoflurane anesthesia (0.7-0.9%) throughout a scan session (Fujimoto et al., 2022). In each session, functional scans were collected before (pre-injection) and after (post-injection) intravenous administration of either vehicle (VEH) or deschloroclozapine (DCZ) 0.1 mg/kg (Nagai et al., 2020). Scans collected during the pre-injection period served as within-session baseline data, allowing us to determine the immediate within-session effects of DREADD activation on rs-FC.

We first conducted a region of interest (ROI) analysis to examine the effect of chemogenetic inhibition of neurons in the amygdala on changes in rs-FC with this area. We used a predetermined anatomy-based ROI for the amygdala that comprised all subnuclei, from the D99 atlas (Saleem et al., 2021). We then separately computed the average correlation between this ROI and every other voxel in the brain collected during the pre-injection and post-injection whole brain resting-state scans. **Figure 3A** shows the rs-FC correlation map from a representative pre-injection scan. In all our pre-injection scans, we found that the amygdala ROIs consistently exhibited strong FC (p<0.05) with the medial prefrontal cortex, ventrolateral prefrontal cortex (vlPFC), and temporal lobes (**Supplemental Figure 1**). This mirrors prior findings (Folloni et al., 2019; Kovacs-Balint et al., 2019) and known anatomical connectivity (Carmichael and Price, 1995; Price, 2003).

**Figure 3.**
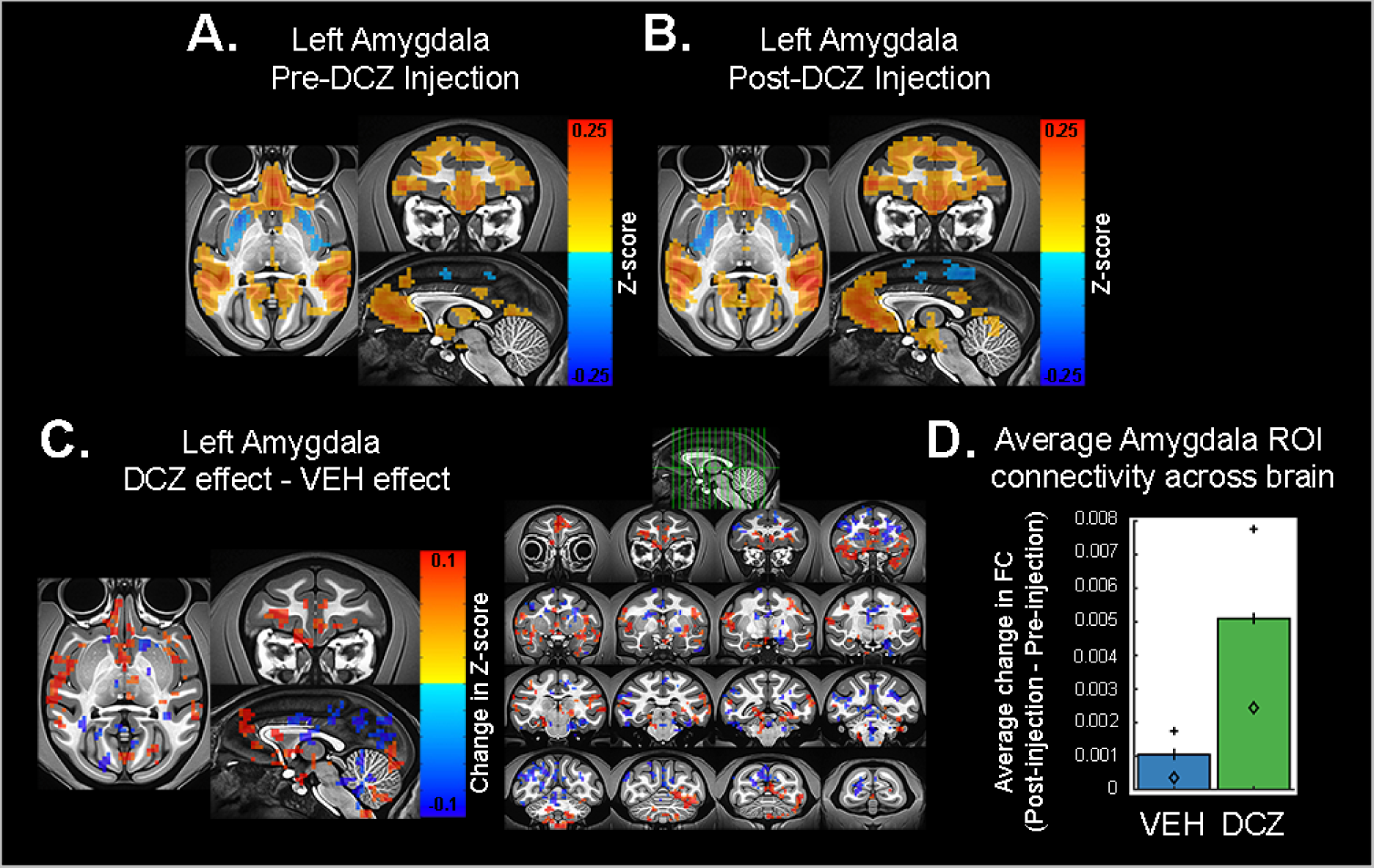
Representative changes in functional connectivity between amygdala and frontal cortex. Changes in FC with a left amygdala seed region in animal L, averaged across two sessions of DCZ testing. Pre-DCZ injection period **(A)** and post- injection period **(B)**. Scale bar indicates z-score of rs-FC. Threshold p=0.0028, cluster size ≥30 voxels, voxel faces touching. **C)** Difference in FC produced by DREADD inhibition [(DCZ post-injection – pre-injection) – (VEH post-injection – pre-injection)], calculated from averaged DCZ post-injection – pre-injection data and averaged VEH post-injection – pre-injection data. Scale bar indicates difference in z-score of rs-FC. Threshold p=0.0485, cluster size ≥30 voxels, voxel faces touching. **D)** Average change in rs-FC between amygdala (bilateral ROI) and all other voxels in the brain after treatment with vehicle or DREADD activation via DCZ. Symbols indicate values for each animal, averaged across sessions. Animal H, diamond; animal L, plus sign. Multiway ANOVA, main effect of drug F[1, 226780] = 272.20, p<0.0001, main effect of animal F[1, 226780] = 186.82, p<0.0001, interaction of drug and animal F[1, 226780] = 64.26, p<0.0001.

To determine changes in rs-FC that specifically occurred as a result of hM4Di-mediated inhibition, we first subtracted the amygdala ROI correlation map (see above) of rs-FC in scans collected pre-injection of VEH or DCZ from those collected post-injection. **Figure 3B** shows an example of the rs-FC correlation map of post-injection data after treatment with DCZ, demonstrating the effects of activating the inhibitory DREADD receptors on the rs-FC of the amygdala. Subtracting the pre-injection data from the post-injection data isolated the change in rs-FC resulting from the injection of VEH or DCZ, as it reduces the influence of session-to-session scanner variability (Fujimoto et al., 2022). Note that this prior work has shown that DCZ alone (at the dose level used here) does not alter rs-FC. We then assessed the effect of DCZ compared to vehicle treatment by subtracting the rs-FC change after vehicle was injected from the rs-FC change after DCZ was injected ([DCZ (post-injection – pre-injection)] – [vehicle (post-injection – pre- injection)]). The results of this subtraction for one animal are shown in **Figure 3C** (more examples shown in **Supplemental Figure 2**). We found that there was a change in rs- FC between the amygdala and the rest of the brain (p<0.0001, **Figure 3D**). Our data show that activating the inhibitory-hM4Di DREADD receptors in amygdala caused an increase in rs-FC between amygdala and multiple cortical areas.

As noted earlier, the amygdala, including the basal nucleus, projects to both cortical and subcortical structures (Amaral et al., 1992; Price, 2003). Further, previous work has shown that inhibition or lesion of one region can result in widespread changes in network connectivity, even across regions that lack anatomical connectivity to the target (Carrera and Tononi, 2014; Vancraeyenest et al., 2020). Thus, we next sought to dissociate the effect of DREADD-mediated amygdala inhibition on global rs-FC connectomes across: 1) the whole brain, 2) cortex, and 3) subcortical areas. To do this, we assessed rs-FC changes using 3 sets of ROIs extracted from the whole brain atlas (D99), a cortical hierarchical atlas (CHARM, Level 3), and a subcortical hierarchical atlas (SARM, Level 3; Seidlitz et al., 2018; Hartig et al., 2021; Jung et al., 2021; Saleem et al., 2021). Importantly, these analyses do not describe the functional connectivity of the amygdala with these regions, but rather generate a connectome in which every ROI’s correlation to every other ROI is measured and the average calculated. This connectome depicts the changes produced in large-scale, brain-wide connectivity as a result of perturbing local activity. It is possible, therefore, that the observed amygdala-frontal cortex rs-FC changes may be influenced by these large-scale network changes.

These connectome analyses revealed that on average activation of the inhibitory DREADD receptors caused a pervasive effect on rs-FC across many brain areas irrespective of whether they were cortical or subcortical (**Figure 4**). Specifically, injection of DCZ was associated with an increase in the rs-FC across the whole brain (D99, **Figure 4A**), a finding that held true when examining changes in rs-FC restricted to only cortical (CHARM, **Figure 4B**) or subcortical structures (SARM, **Figure 4C**). Notably, there were differences between the subjects in each of the three analyses, with a significant interaction between subject and treatment for the whole brain and subcortical rs-FC analyses only. In the analysis of rs-FC among subcortical structures the two animals had divergent patterns; activating the inhibitory DREADD receptors causes an increase on average in rs-FC in animal H, whereas animal L showed a decrease on average (**Figure 4C**, symbols). Such a difference may be related to differences in expression of the DREADD receptors in amygdala (**Figure 2**; see discussion below). Overall, these analyses reveal that DREADD-mediated inhibition of neurons in the amygdala is associated with an increase in rs-FC both within specific circuits between the amygdala and cortical regions, and across networks spanning the whole brain, with the most consistent results seen in cortical networks.

**Figure 4.**
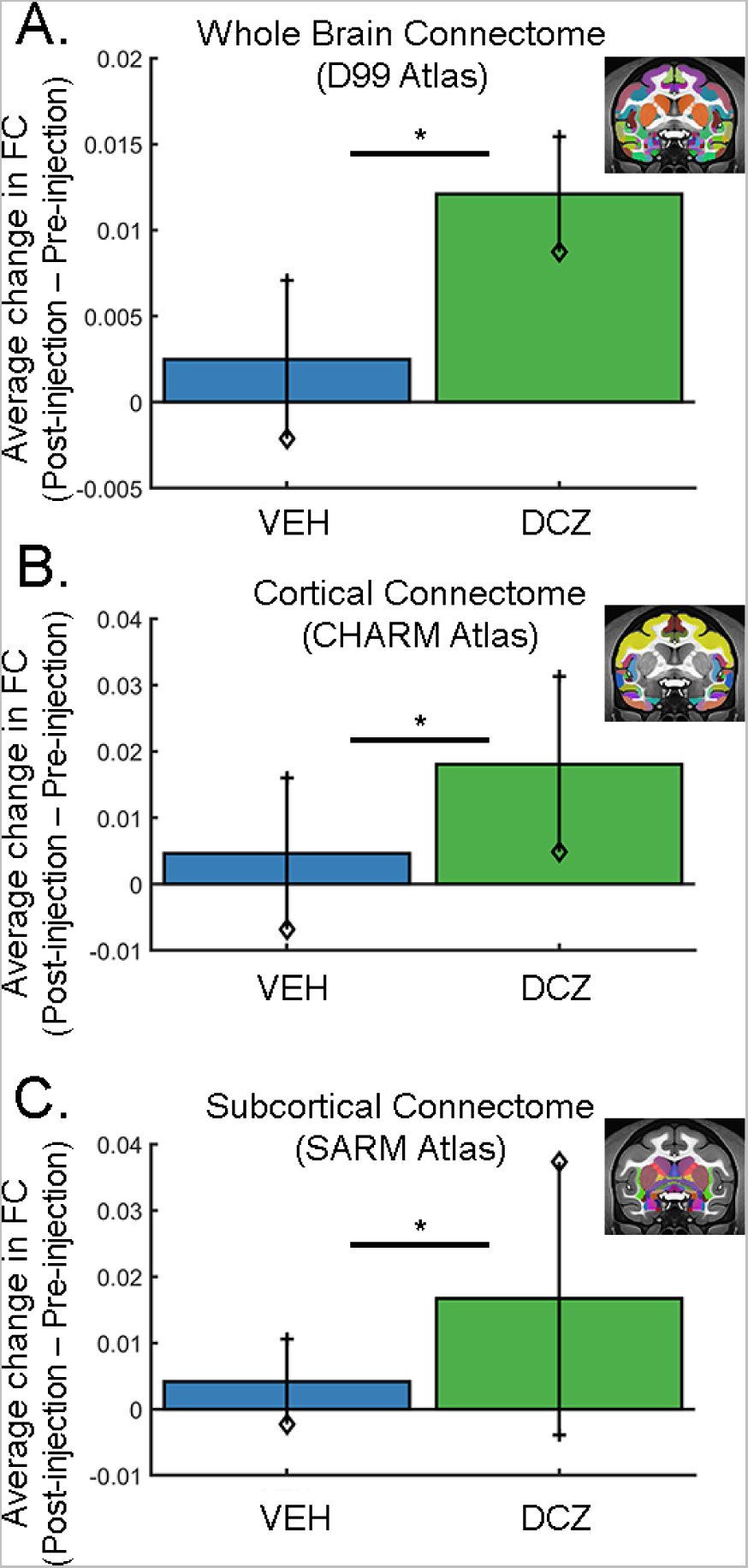
Whole brain fMRI functional connectome altered by chemogenetic inhibition of amygdala. Average FC across all ROIs calculated across a standard whole brain atlas (**A**), a cortical atlas (**B**) and a subcortical (**C**) atlas. Activation of DREADDs via DCZ increased global rs-FC across all three networks. Symbols denote average difference in rs-FC for each animal, averaged across sessions. Animal H, diamond; animal L, plus sign. Error bars represent SEM. Atlas image insets shown on NMT v2 (Seidlitz et al. 2018, Hartig et al. 2021, Jung et al. 2021, Saleem et al. 2021). Multi-way ANOVA analyses: Whole brain, main effect of drug F[1, 146884] = 728.35, p<0.0001 and subject F[1, 146884] = 498.91, p<0.0001, interaction between subject and drug F[1, 146884] = 12.54, p=0.0004. Cortical atlas, main effect of drug F[1, 5036] = 34.71, p<0.0001 and subject F[1, 5036] = 115.59, p<0.0001. Subcortical atlas, main effect of drug F[1, 4484] = 44.61, p<0.0001 and subject F[1, 4484] = 56.76, p<0.0001, interaction between subject and drug F[1, 4484] = 206.47, p<0.0001.

### DREADD-mediated changes in local circuit activity and LFP coherency

There are two possible reasons that the inhibitory DREADD receptors in the amygdala produced an increase in fMRI rs-FC. One possibility is that activating the inhibitory DREADD receptors may reduce local neural activity, causing an attenuation of high frequency activity. Consequently, slow oscillations would dominate both local neural activity and the fMRI signals, resulting in more temporally correlated activity between distributed brain areas. Such a pattern would be consistent with work in rodents (Rocchi et al., 2022). Alternatively, it is possible that activating the inhibitory DREADD receptors actually causes an *increase* in neural activity in amygdala through local recurrent mechanisms, such as the release of excitatory neurons from lateral inhibition and/or inhibition of inhibitory interneurons. The observed increase in rs-FC would therefore be the result of this increased activity driving long-range functional interactions in the brain. To determine the relationship between fMRI rs-FC, which represents an indirect measure of correlated neural activity, and direct measures of neural activity, we conducted extracellular neural recordings in our resting-state paradigm in the same animals (**Figure 1C**).

Both animals underwent electrophysiology recordings using 2 or 3 16-channel linear arrays simultaneously targeting the amygdala and vlPFC (**Figure 5A**). We focused on the relationship between amygdala and the vlPFC (area 12, located laterally to the lateral orbital sulcus on the ventral surface of the brain) because: 1) the amygdala sends direct mono-synaptic input to this area (Carmichael and Price, 1995); 2) this pathway is functionally engaged during reward learning (Chau et al., 2015); and 3) we found that rs- FC was increased between amygdala and vlPFC when the DREADDs were activated (**Figure 3**). The recording sites in amygdala and vlPFC were therefore determined based on the prior fMRI analyses and matched to the post-mortem histology (**Figure 2**). In order to compare the results across modalities, neurophysiology data were acquired under conditions identical to those used to acquire the fMRI data (i.e. under light isoflurane anesthesia, Fujimoto et al., 2022). We investigated changes in both spiking activity and LFP signal.

**Figure 5.**
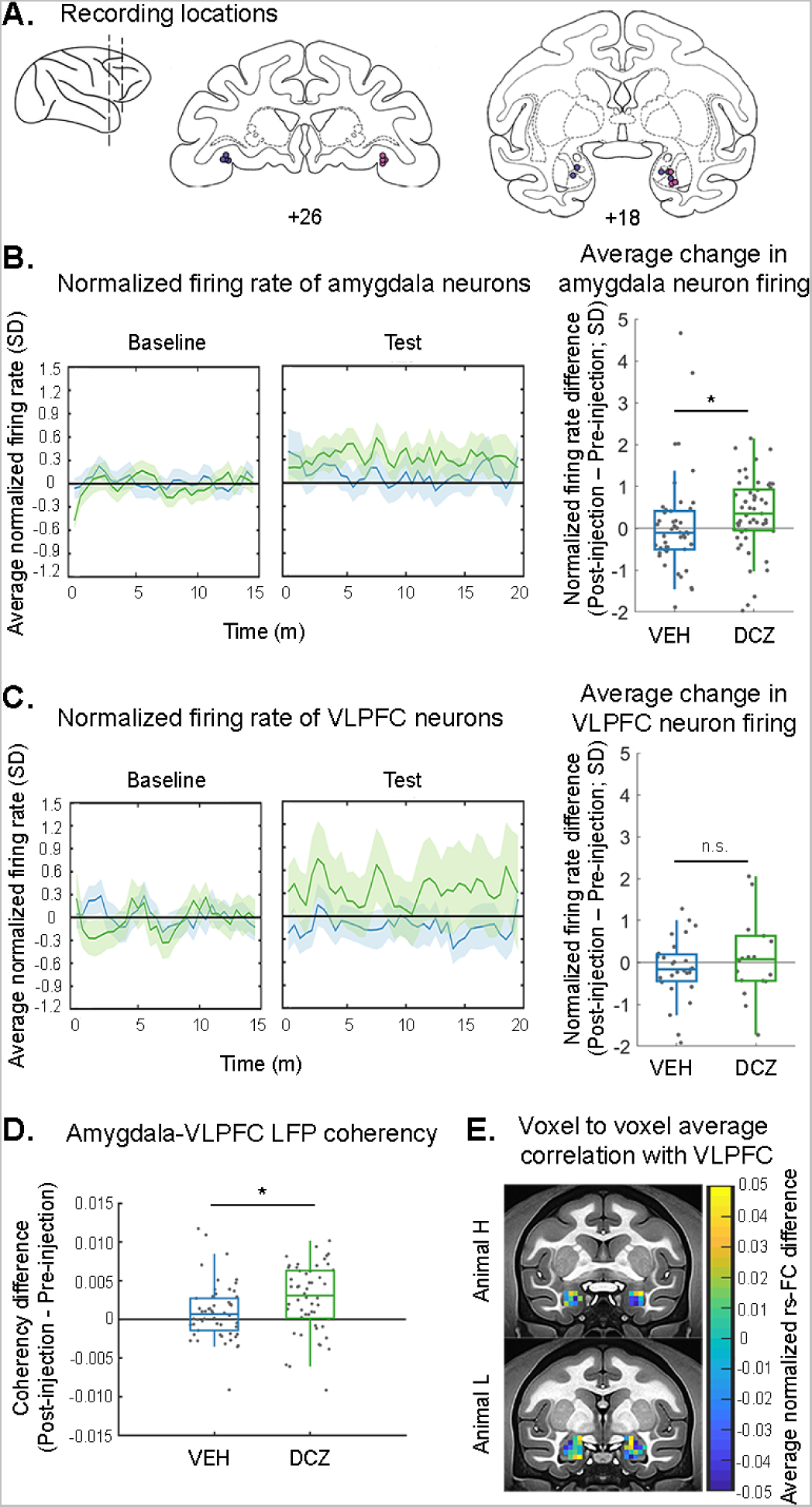
Inhibition of amygdala produces increased functional connectivity as measured by LFP coherency and rs- fMRI, as well as increased neural spiking. **A)** Recording sites in vlPFC and amygdala, validated by histology. **B-C)** Average (±SEM) time course (left) and summary data (right) of neural spiking activity in the amygdala (**B**) or vlPFC (**C**) after treatment with DREADD actuator DCZ or control. For visualization purpose, the average normalized firing rate change over time was smoothed using a moving average window of 2sec (4 bins). Only amygdala neurons show an increase in average firing rate after chemogenetic inhibition of amygdala via DCZ (Wilcoxon signed-rank test; p=0.002. Amygdala VEH p=0.46; vlFPC DCZ p=0.68, vlPFC VEH p=0.29). There was a significant difference between amygdala neuron firing rate after treatment with DCZ as compared to VEH (Kruskall-Wallis; Χ^2^ [1,97, N = 99] = 7.3, p = 0.0069), but no difference due to treatment in the vlPFC (Χ^2^ [1,43, N = 45] = 0.48, p = 0.49). Single neurons denoted by dots. **D)** Average (±SEM) change in LFP coherency at each amygdala electrode site pair with all vlPFC electrode sites. Dots indicate amygdala contact pairs. Multiway ANOVA, main effect of drug F[1, 102] = 5.64, p=0.02 **E)** Average change in FC between each amygdala ROI voxel with all vlPFC (area 12o/l) voxels, shown separately for each animal. Right and left rs-FC maps were calculated separately.

First, we analyzed neuronal spiking activity in both recorded areas to establish whether the observed rs-FC changes were directly associated with changes in firing rate. We recorded the activity of 99 single neurons in amygdala across the pre- and post-injection periods for DCZ or VEH treatment (see **Table 1**). We found that activating the inhibitory DREADD receptors with DCZ increased the firing rate of the recorded amygdala neurons as compared to VEH treatment (**Figure 5B**). Additionally, only amygdala neurons after DCZ treatment altered their firing rate significantly as compared to the pre-injection period. In the vlPFC, DCZ did not alter firing rate as compared to VEH, and neither treatment altered firing rate significantly from the pre-injection period (**Figure 5C**). These data indicate that activation of the inhibitory DREADD receptors in amygdala was associated with an increase in neural activity locally, but not at distant locations such as vlPFC. This confirms that the changes in activity we observed were not simply the result of off-target effects of DCZ.

**Table 1.** Summary of recorded neurons.

| **Table 1. Summary of recorded neurons** |  |  |  |  |  |  |
| --- | --- | --- | --- | --- | --- | --- |
|  | **VEH** |  | **DCZ** |  | **CNO** |  |
|  | **AMG** | **vIPFC** | **AMG** | **vIPFC** | **AMG** | **vIPFC** |
| **Mk L** | 28 | 8 | 31 | 6 | 21 | 7 |
| **Mk H** | 18 | 19 | 22 | 12 | 7 | 8 |
| **Total** | **46** | **27** | **53** | **18** | **28** | **15** |

Next, we looked at the relationship between LFP activity in our regions of interest by extracting the LFP coherency between amygdala and vlPFC. We first averaged coherograms across contacts, sessions and periods, in order to reveal in an unbiased manner a frequency band of interest for each animal, i.e. a band in which we observed a clear peak in the coherency. For both animals, a peak in coherency was evident from 6.5 to 14.5 Hz, a range centered on the alpha band (**Supplemental Figure 3A**). We then extracted the average alpha coherency of each amygdala contact with all vlPFC contacts and subtracted the pre-injection average coherency value from the post-injection average coherency value. Similar to the normalization procedure conducted on the rs-FC data, this approach reduced the impact of cross-session variability in LFP coherency signals. We observed a significant increase in alpha-band coherency caused by DCZ-mediated activation of the DREADD receptors (**Figure 5D**). We did not observe any changes in power across this frequency band after treatment with DCZ as compared to VEH (**Supplemental Figure 3B**). This result suggests that the DREADD-mediated changes in rs-FC that we observed between amygdala and vlPFC were related to increased LFP coherency in the alpha band, in keeping with some prior reports (for example, Wang et al., 2012).

### Comparison of fMRI rs-FC with vlPFC across the spatial extent of the amygdala

The previous analyses indicated that rs-FC between amygdala and vlPFC is related to LFP coherency in the alpha band. We focused our extracellular recordings on the basal nucleus of the amygdala, where we had originally targeted for DREADD transfection surgery, and on Walker’s area 12o of the vlPFC, a region known to receive anatomical projections from the basal nucleus. To compare these results to our fMRI rs- FC findings with more spatial specificity, we conducted an additional fine grain analysis specifically looking at rs-FC between amygdala and vlPFC. We also conducted this analysis to see if there were qualitatively similar effects on rs-FC and the known expression patterns of DREADD receptors in each subject. Thus, these analyses were conducted separately for each animal.

We first calculated the mean connectivity of each amygdala voxel with the vlPFC. Here functional connectivity was calculated between each voxel in the vlPFC ROI and each voxel in the amygdala ROI in both the pre-injection and post-injection periods for either DCZ or vehicle. The resulting pre-injection connectivity matrices were subtracted from the post-injection connectivity matrices, and then the VEH matrices were subtracted from the DCZ matrices ([DCZ post-injection – pre-injection] – [VEH post-injection – pre-injection]). The correlation values at each amygdala voxel were then averaged such that the resulting values reflect each amygdala voxel’s average change in rs-FC with all vlPFC voxels following DCZ administration compared to vehicle. We then projected this functional connectivity matrix back onto the amygdala and reasoned that with higher DREADD expression, a subregion of amygdala should exhibit stronger changes in rs-FC with vlPFC (**Figure 5E**).

In animal H, dorsal amygdala regions across the medial-lateral extent showed stronger positive rs-FC changes with vlPFC after DREADD activation, while ventral regions showed overall negative rs-FC changes with vlPFC. This pattern of changes appears to align closely with the patterns of DREADD receptor expression that we observed in this animal (**Figure 2**). In animal L, medial amygdala voxels across the dorsal-ventral extent, as well as voxels in ventrolateral amygdala, showed a positive rs-FC change with vlPFC. Again, the spatial pattern of rs-FC effects appears to mirror the spatial pattern of DREADD receptor expression in this subject’s amygdala. In summary, the direction of voxel-based rs-FC between amygdala and vlPFC was relatively heterogenous, suggesting mixed effects at the level of local neuronal circuitry. The strongest positive changes in rs-FC appeared centered on the site of the basal nucleus in both animals, in agreement with our LFP coherency results showing increased coherency between this region of the amygdala and vlPFC.

### Activation of inhibitory DREADDs with clozapine-n-oxide does not reproduce the effects obtained with DCZ

Throughout this study, we used DCZ, a recently validated compound (Nagai et al., 2020) to activate the inhibitory DREADD receptors. This actuator is highly selective for DREADD receptors and enables precision targeting without concern for off-target effects on behavior or whole-brain functional connectivity (Upright and Baxter, 2020; Fujimoto et al., 2022; Cushnie et al., 2023). However, much of the initial and current research incorporating DREADDs uses clozapine-N-oxide (CNO; Armbruster et al., 2007; Grayson et al., 2016). While CNO has a high affinity for the DREADD receptors, it has been found to have low blood brain barrier penetrance and is back-metabolized to clozapine, causing non-specific effects as clozapine binds to and activates endogenous receptors in the brain (Gomez et al., 2017). As CNO continues to be used in basic neuroscience, and because CNO was used in a prior study on the effects of DREADD-mediated inhibition of the amygdala on rs-FC (Grayson et al., 2016), we collected additional fMRI and neurophysiology data where CNO was injected instead of DCZ.

Our seed-based fMRI analysis produced an inconsistent pattern of rs-FC between frontal cortex and amygdala such that some regions showed increased rs-FC and others decreased (**Figure 6A**). Throughout the brain, treatment with CNO decreased rs-FC with the amygdala (p<0.0001, **Figure 6B**). Similarly, the global, atlas-based rs-FC analysis showed, in contrast to our experiments using DCZ, a significant decrease in overall rs- FC when the DREADD receptors were activated by CNO (**Figure 6C**). This effect was seen as well across subcortical structures, but not across cortical areas.

**Figure 6.**
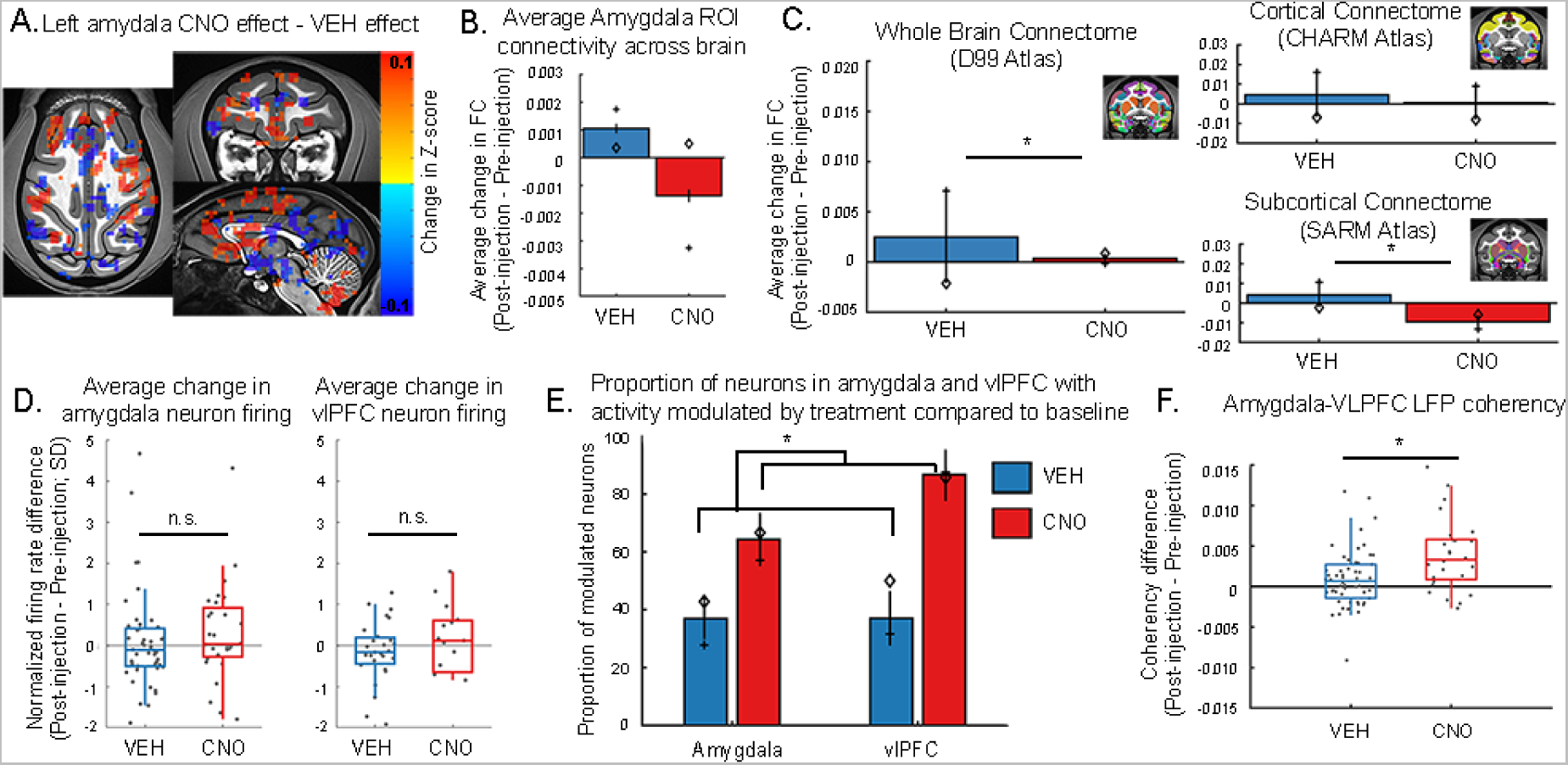
Activation of inhibitory DREADDs by less specific actuator CNO compared to DCZ produces similar changes in frontal-limbic functional connectivity but differently affects large scale connectomes. **A)** Difference in FC with left amygdala seed region in animal L produced by DREADD inhibition with CNO (as in Figure 3C). Scale bar indicates difference in z-score. Threshold p=0.0485, cluster size ≥30 voxels, voxel faces touching. **B)** Average change in rs-FC between amygdala (bilateral ROI) and all other voxels in the brain after treatment with vehicle or DREADD activation via CNO. Symbols indicate values for each animal (averaged across VEH sessions). Animal H, diamond; animal L, plus sign. Multiway ANOVA, main effect of drug F[1, 170084] = 67.70, p<0.0001, main effect of animal F[1, 170084] = 16.23, p<0.0001, interaction of drug and animal F[1, 170084] = 76.22, p<0.0001. **C)** Average (±SEM) rs-FC across all ROIs calculated using a standard whole brain atlas, as well as specifically within cortical and subcortical regions. Compared to vehicle, CNO decreased global rs-FC across the whole brain, and specifically across subcortical regions. Whole brain multiway ANOVA, main effect of drug (F[1, 110012] = 24.49, p<0.0001), subject F[1, 110012] = 90.89, p<0.0001, interaction between subject and drug F[1, 110012] = 140.66, p<0.0001. Cortical atlas multiway ANOVA, main effect of subject F[1, 3776] = 51.04, p<0.0001, no main effect of drug F[1, 3776] = 2.17, p=0.14, no interaction F[1, 3776] = 1.05, p=0.31. Subcortical atlas, main effect of drug (F[1, 3362] = 42.99, p<0.0001), interaction between subject and drug F[1, 3362] = 23.28, p<0.0001, no main effect of subject F[1, 3362] = 1.77, p=0.18. **D)** Amygdala DREADD inhibition via CNO did not alter neural spiking activity in the amygdala (left; Kruskall-Wallis; Χ^2^ [1,72, N = 74] = 2.0, p = 0.16) or vlPFC (right; Kruskall-Wallis; Χ^2^ [1,40, N = 42] = 0.52, p = 0.47) as compared to vehicle. In no condition was the firing rate significantly altered compared to pre-injection (Wilcoxon signed-rank test; Amygdala CNO p=0.33; amygdala VEH p=0.46; vlFPC CNO p=0.85, vlPFC VEH p=0.29). However, CNO treatment did **E)** significantly increase the proportion of modulated neurons across the amygdala and vlPFC. Dots indicate single neurons. Error bars indicate SEM. Linear mixed-effects model (T[1,112] = 2.25, p=0.03). **F)** DREADD activation with CNO significantly increased LFP coherence between amygdala and vlPFC. Average (±SEM) change in LFP coherency at each amygdala electrode site pair with all vlPFC electrode sites. Dots indicate amygdala contact pairs. Multiway ANOVA, main effect of treatment (F[1, 78] = 10.73, p=0.002).

Using multi-contact linear arrays, we recorded the activity of 28 neurons in amygdala and 15 neurons in vlPFC, as well as LFPs from both areas, before and after intravenous injection of CNO. We did not observe significant changes in neural spiking after CNO injection (**Figure 6D**). Despite this, CNO appeared to be associated with more heterogeneous changes in firing rates. CNO significantly modulated a larger proportion of neurons across amygdala and vlPFC areas as compared to vehicle (**Figure 6E**). These results suggest that CNO is modulating neural spiking in our target regions. However, it is producing both increases and decreases in neural spiking to the extent that the overall population activity is not altered. In the frequency band centered on alpha (6.5-14 Hz; described above) we observed an increase in coherency between amygdala and vlPFC following CNO injection (**Figure 6F**). This effect was similar to what we observed when DCZ was administered (**Figure 5**). Taken together, these results suggest while both DCZ and CNO are causing activation of DREADD receptors, CNO’s off target effects cause broad changes in neural circuits and lead to the opposite pattern of rs-FC and neural activity compared to DCZ. These findings further highlight the use of DCZ as an actuator drug for DREADDs and reinforce the importance of controlling for off-target effects when examining changes in rs-FC.

## DISCUSSION

We investigated the neural correlates underlying fMRI rs-FC in fronto-limbic circuits. Specifically, we utilized a chemogenetic approach in macaque monkeys to transiently alter the activity of amygdala neurons and assessed the effects on fMRI rs-FC and neural activity markers. We found that activating the inhibitory DREADD receptors was associated with an increase in rs-FC between amygdala and regions that are functionally and anatomically connected to amygdala such as frontal and temporal cortex. We also observed a general increase in functional connectivity across the brain, most notably across cortical regions. When we probed the neurophysiological basis of these effects, we found that chemogenetic inhibition of amygdala neurons produced a modest increase in the firing rate of amygdala neurons. We also characterized the changes in neurophysiological markers of rs-FC between the amygdala and vlPFC, a region known to receive direct projections from the amygdala. We found that LFP coherence in the alpha band between these brain areas also increased after chemogenetic inhibition of the amygdala, similar to the increase in fMRI rs-FC. This pattern of effects provides convergent evidence that coherency in the LFP in low frequency may underlie rs-FC measures in frontal-limbic circuits.

### Basis of resting-state functional connectivity in frontal and limbic circuits

Prior work comparing hemodynamic signals to direct recordings of neuronal activity has reported that fMRI rs-FC between areas is closely related to patterns of synchronous or coherent activity in specific frequency bands of the LFP (Lurie et al., 2020). However, the specific frequency band that underlies rs-FC in the brain has been harder to pinpoint. For example, in humans rs-FC between parts of lateral and medial parietal cortex is related to correlations in the low beta and high-gamma frequency bands recorded using electrocorticography (Foster et al., 2015). By contrast, LFP coherence in the alpha band across macaque pulvinar, lateral intraparietal area, V4, and TEO has been shown to correlate most strongly with fMRI rs-FC (Wang et al., 2012). Other studies have similarly highlighted the importance of other LFP frequency bands for rs-FC (Lu and Haber, 1992; Hacker et al., 2017). Our experiments revealed that LFP coherency was strongest centered on the alpha band (between 6.5-14.5 Hz) in amygdala to frontal cortex and thus appears to agree with the findings of Wang and colleagues that looked at cortico-cortical rs-FC in macaques (Wang et al., 2012). This should not, however, be taken as evidence that coherence in the alpha band is the only mechanism underlying all rs-FC in the brain. A number of groups have shown that the frequencies that contribute to rs-FC vary across the brain depending on the specific circuit or functional network (Siegel et al., 2012; Hipp and Siegel, 2015; Hacker et al., 2017). What we reveal here is that in pathways connecting the amygdala to frontal cortex, that rs-FC is associated with alpha band coherence, knowledge that is likely important for developing targeted therapies that aim to alter functional networks, a point we take up below.

As we noted above, the majority of prior work on the basis of rs-FC specifically looked at the correlation between fMRI and neural activity without determining whether perturbing activity in a circuit would cause a corresponding change in both rs-FC and its underlying neural mechanisms. Doing so would provide evidence for the neural mechanism of rs-FC. Prior work using lesions in macaques revealed that sectioning the corpus callosum was associated with a decrease in intra-hemispheric rs-FC (O’Reilly et al., 2013). This work and others using lesions (Chen et al., 2015; Adam et al., 2020) were, however, unable to reveal the associated changes in neural activity that followed the lesion-induced changes in rs-FC. As lesions involve local cell death, these studies also cannot assess how increases in local activity might change rs-FC. Aspiration lesions have off-target effects (Meunier et al., 1999; Rudebeck et al., 2013b), further limiting the insights that can be gained from this approach. To overcome these issues, we used DREADDs to specifically target neurons in amygdala. Previous work supports the use of this approach in macaques for studying changes in rs-FC (Grayson et al., 2016; Vancraeyenest et al., 2020; Hirabayashi et al., 2021). By successfully combining a chemogenetic approach with fMRI and neurophysiology, our results therefore provide causal evidence that increasing rs-FC between amygdala and vlPFC is directly related to increased synaptic modulation, specifically in the alpha band of the LFP. Thus, our causal approach extends prior work by revealing the relationship between rs-FC and coherency in neural activity.

Given the similarity between our findings and those of Wang and colleagues (Wang et al., 2012) it is likely that alpha band coherence underlies rs-FC in other fronto-limbic or cortico-cortical pathways. Despite this, the effects of chemogenetic modulation of rs-FC and neural activity still are likely to be specific to the circuits under investigation. This difference between neural circuits was highlighted by a recent study in rodents that also used DREADDs to manipulate rs-FC. Here the authors found a similar relationship between LFP coherency and rs-FC (Rocchi et al., 2022). Specifically, chemogenetically inactivating prelimbic cortex led to an increase in rs-FC and LFP coherency between prelimbic cortex and thalamus. Unlike our findings, LFP coherency was increased in the delta band (1-4 Hz). Thus, both ours and the study by Rocchi and colleagues consistently show that rs-FC is directly and causally related to LFP coherence. On the other hand, there are clear differences between the effect of inhibitory DREADDs on neural activity and the specific LFP frequency underlying rs-FC between the two studies (delta vs alpha). It is likely that such variation in findings is related to the circuits under investigation (Hipp and Siegel, 2015), the species being studied (mice versus macaques), or some combination of both.

It is important to note that previous chemogenetic studies in macaques, including our own, were not able to determine if the recorded neurons are those that have expressed DREADD receptors, or if they are simply communicating with DREADD-expressing neurons. Only three recent studies in macaques have recorded spiking activity from neurons after DREADD transfection in the target region (Deffains et al., 2021; Hasegawa et al., 2022; Perez et al., 2022), and all observed a decrease in firing rate after chemogenetic inhibition of the target. However, none of these studies recorded from the amygdala, or targeted fronto-limbic circuitry. Indeed, our finding that activating an inhibitory DREADD in macaque amygdala led to increased spiking in this area (**Figure 5**) could be related to the composition of this structure in primates. Recent studies have shown that there is an increased diversity of interneurons in macaque amygdala compared to rodents (Beyeler and Dabrowska, 2020; McDonald and Augustine, 2020; McDonald, 2021). Thus, we speculate that inhibiting these cells may have caused a paradoxical disinhibition of local amygdala circuits, which led to increased extracellular activity. A second possibility is that the inhibition of local excitatory neurons that drive inhibitory networks within the amygdala could have led to increased local activity (Smith et al., 2000). These explanations are not mutually exclusive, and future work may serve to clarify the mechanisms at play. In summary, our findings and those of Rocchi and colleagues emphasize the importance of confirming the effect of chemogenetic modulation on neural activity. Further, they also highlight the power of chemogenetic manipulation for revealing the basis of rs-FC.

### The differential effect of DREADD actuators on fMRI and neural markers of functional connectivity

To date, few studies have investigated the neural basis of functional connectivity in limbic circuits, either generally or in specific relation to the amygdala. A previous paper examining changes in fMRI rs-FC after DREADD-mediated inhibition of the amygdala found decreased FC between amygdala and frontal regions after administration of the ligand CNO (Grayson et al., 2016). On the face of it, this result appears to be counter to what we report here, namely that DREADD-mediated inhibition of amygdala neurons increased rs-FC. In the years since Grayson et al. was published, refinements have been made in macaque anesthetized fMRI protocols (Xu et al., 2019; Autio et al., 2021; Fujimoto et al., 2022), DREADD constructs (Galvan et al., 2019), and DREADD actuators (Gomez et al., 2017; Nagai et al., 2020).

In part because of these refinements, we collected our data while animals were maintained under light (≤0.9%) isoflurane, allowing for maximal preservation of resting state network activity (Hutchison et al., 2014; Wu et al., 2016; Areshenkoff et al., 2021; Autio et al., 2021); we consider this particularly important given that cortical rs-FC is most strongly impacted by increasing doses of anesthesia (Hutchison et al., 2014; Lv et al., 2016; Giacometti et al., 2022). In light of work finding that DREADD constructs carrying an mCherry fusion protein are only sparsely expressed at the cell surface (Galvan et al., 2019), we chose to use a DREADD construct that instead contains an HA tag. Finally, we used a highly selective, potent DREADD actuator developed for use in macaques, DCZ, rather than CNO (Nagai et al., 2020). The latter has been shown to be metabolized to clozapine, which has broad affinity for endogenous neuromodulatory receptors in the brain in addition to DREADDs (Gomez et al., 2017).

Although the anesthesia dose and choice of viral vector used for DREADD transfection we used both likely impacted our results, we reasoned that the diminished effects of DREADDs and off-target effects caused by CNO/clozapine might be the key difference between the fMRI rs-FC experiments in Grayson’s study and our own. Thus, we conducted one session of testing in each animal and each experimental paradigm in which we administered CNO, rather than DCZ. As our results show, administration of CNO, as compared to DCZ, in our subjects caused a pattern of effects that was more similar to what Grayson and colleagues reported (**Figure 6**). What is notable is that administration of either CNO or DCZ caused the same increase in LFP coherency between amygdala and vlPFC. Thus, our data appear to show that activating inhibitory DREADD receptors, regardless of the actuator used, has a consistent effect on neurophysiological markers of rs-FC, but conversely that the choice of actuator has profoundly different effects on fMRI rs-FC (Nagai et al., 2020; Fujimoto et al., 2022).

Why CNO activation of DREADD receptors causes such a different pattern of rs- FC is an open question, but likely relates to the fact that this ligand is metabolized to clozapine (Gomez et al., 2017). Clozapine has high affinity for D1 and D2 dopamine and 5-HT2A and 5-HT2C serotonin receptors (Phillips et al., 1994) that are widely distributed across the brain. It is likely that the off-target effects of clozapine on these receptors underlies the wide-scale effects of DREADD inactivation of amygdala on rs-FC.

### Conclusions and relevance to psychiatric disorders

Our multidisciplinary approach to establishing the basis of rs-FC in fronto-limbic circuits provides new insight into the mechanisms underlying this increasingly used and clinically useful measure of functional interactions in the brain. In recent years, patterns of rs-FC in fronto-limbic circuits are being explored as potential biomarkers of different psychiatric disorders (Parkes et al., 2020). With the advent of large cohort imaging studies, it has even been possible to identify different subtypes of depression (Chai et al., 2023), autism (Ren et al., 2023), anxiety (Li et al., 2023), and schizophrenia (Zhu et al., 2016) based on their differential patterns of rs-FC. One potential use for these biomarkers is to identify and then target neuromodulation therapies such as deep brain stimulation (DBS) to specific circuits to ameliorate symptoms (Holtzheimer and Mayberg, 2011). Thus, our finding that rs-FC in a fronto-limbic circuit is directly related to coherence in the alpha band of the LFP is translationally relevant as it establishes the baseline for patterns of rs- FC in healthy circuits that are often implicated in psychiatric disorders (Klein-Flügge et al., 2022). Consequently, this information could be used to optimize treatment approaches such as DBS to attempt to heighten alpha band communication between affected areas.

## METHODS AND MATERIALS

### Subjects

Two male adult rhesus macaques (*Macaca mulatta*), age 7, were used as subjects in this study. Both animals (animal H, animal L) underwent anesthetized functional MRI scans before and after DREADD transfection surgery. Both animals also underwent anesthetized electrophysiological recording from the amygdala and frontal cortex after DREADD transfection surgery. All procedures were reviewed and approved by the Icahn School of Medicine Animal Care and Use Committee.

### Surgical targeting MRI

A pre-operative MRI scan was collected in each animal to target the amygdala. The animal was sedated with a mixture of ketamine (5 mg/kg) and dexmedetomidine (0.0125mg/kg), intubated, and placed into a standard MRI-compatible stereotaxic frame. Animals were then transported to the imaging facility where they were maintained at a stable plane of anesthesia using isoflurane (1%–4%) during image acquisition. Between 2-4 3D T1-weighted images (0.5 mm isotropic, TR/TE 2500/2.81 ms, flip angle 8°) were acquired on a Siemens MAGNETOM 3 Tesla Skyra scanner using a custom-built 4- channel phased array surface coil with local transmit function (Windmiller-Kolster Scientific, CA).

### Surgical Procedures

All surgical procedures were conducted under isoflurane (2-3%) general anesthesia.

First, all animals underwent an AAV vector injection surgery. AAV vector encoding for inhibitory DREADD (AAV5-SYN1-hM4Di-HA 1.7*10^13^ GC/ml (Addgene, Watertown MA)) was injected into the bilateral basolateral amygdala stereotaxically. Six sites within the amygdala were chosen to achieve full coverage of the structure. One site was chosen to target basal amygdala, and a further two were located 2mm posteriorly and targeted basal accessory, basal, and medial nuclei. Each of these sites had an additional site targeted 2mm ventrally. Injection sites were chosen by consulting each animal’s pre-surgical MRI for precise targeting.

To inject the viral vector, first a small craniotomy was made in the skull over each amygdala. Virus was drawn up into a syringe and the surface of the needle was wiped down with alcohol and sterile saline. The syringe was positioned using an arm attachment on the stereotaxic frame, and the needle was lowered into the brain. To reduce both pressure at the injection site and the chance of tissue blockage at the tip of the needle, at each of the three locations the needle was first lowered 2mm ventral to the target, then pulled up to the correct coordinates. The more ventral site was injected first, then the more dorsal site. Injections were made by hand at an approximate rate of 0.2 ul/min. After each injection, at least five minutes elapsed before the needle was moved to reduce any leakage of virus. 3 ul were injected at each site for a total of 18 ul per hemisphere. At least two months elapsed after surgery before any experiments were conducted to allow for full viral expression.

Once neuroimaging experiments were completed, a rectangular acrylic chamber (Rudebeck et al., 2013a) was implanted with ceramic screws (Thomas Recording, Giessen, Germany) and biocompatible acrylic (Lang Dental, Wheeling, IL), which allowed access to both the amygdala and ventrolateral prefrontal cortex. Following a 2-week recovery period, a craniotomy was made inside the chamber. Coordinates for surgical planning were determined with a pre-acquired T1-weighted MR image.

### Drug Preparation

Deschloroclozapine (DCZ) solution was prepared using DCZ powder (HY-42110, MedChemExpress), dissolved in dimethyl sulfoxide (DMSO, 2% of total volume) and then diluted in saline to a concentration of 0.1 mg/kg (Nagai et al., 2020), total volume of 1ml. Clozapine-N-oxide (CNO) solution was prepared using CNO powder (Jin lab, Mount Sinai School of Medicine), dissolved in dimethyl sulfoxide (DMSO, 2% of total volume) and then diluted in saline to a concentration of 10 mg/kg, total volume 10 ml. Vehicle solution was prepared as 2% DMSO diluted in 1ml saline. All solutions were prepared within 30 min of usage.

### fMRI data acquisition

We followed an identical protocol for functional imaging as described in our earlier paper (Fujimoto et al., 2022). Animals were first sedated with ketamine (5mg/kg) and dexmedetomidine (0.0125mg/kg) 1-1.5 h before the data collection to prevent detrimental effects of ketamine on neural activity. Animals were placed inside an MRI-safe plexiglass primate chair in the sphinx position during image acquisition (Rogue Research, Cambridge MA). The animal’s head was not restrained, but supported with towels and flexible bandages to allow for stability with minimal discomfort. This setup allowed for the use of lower maintenance isoflurane levels during image acquisition; isoflurane anesthesia was maintained at 0.7-0.9% throughout all functional imaging sessions. The use of low-level anesthesia allows for preservation of resting-state networks (Hutchison et al., 2013; Wu et al., 2016; Giacometti et al., 2022). To minimize the effect of physiological changes in the neural activity, vital signs (end-tidal CO2, body temperature, blood pressure, capnograph) were continuously monitored and maintained throughout an experimental session.

In each session, pre-injection and post-injection (VEH, DCZ, CNO) functional scans were collected. First, a set of setup scans were acquired, which included shimming based on the acquired fieldmap. A session specific 3D T1-weighted image (0.5 mm isotropic, TR/TE 2500/2.81 ms, flip angle 8°) was acquired. Following monocrystalline iron oxide nanoparticle (MION) injection (i.v.) (Leite et al., 2002; Russ et al., 2021), three functional scans (Echo Planar Images (EPI): 1.6 mm isotropic, TR/TE 2120/16ms, flip angle 45°, 300 volumes per each run) were obtained for pre-injection functional resting states. Next, either vehicle, DCZ (0.1 mg/kg), or CNO (10 mg/kg) was administered (i.v.). In sessions where DCZ or VEH were tested, we allowed 15 mins after the injection to ensure trafficking of the drug to the brain (Nagai et al., 2020), prior to the collection of an additional three functional runs. The order of DCZ and VEH testing was counterbalanced across animals. Both monkeys H and L completed two sessions each of imaging with VEH and DCZ injection. After the conclusion of the VEH and DCZ sessions, both animals completed one CNO session. In this session, the post-injection EPI data were collected ∼30 minutes after CNO injection to allow for the longer time course of CNO trafficking and metabolic processing (Nagai et al., 2020).

### fMRI data analysis

Functional imaging data were preprocessed with standard AFNI/SUMA pipelines (Cox, 1996; Jung et al., 2021; Fujimoto et al., 2022). Raw images were first converted into NIFTI data file format and ordered into BIDS format (Gorgolewski et al., 2016). The T1-weighted images were spatially normalized, then skull-stripped using the U-Net model built from Primate Data-Exchange open datasets (Wang et al., 2021). The normalized skull-stripped T1-weighted image was then aligned to the NMT version 2.0 atlas (Seidlitz et al., 2018), along with subject-specific versions of the D99 (Saleem et al., 2021), Cortical Hierarchical Atlas of the Rhesus Macaque (CHARM) (Jung et al., 2021) and Subcortical Atlas of the Rhesus Macaque (SARM) atlases (Hartig et al., 2021).

The functional data were preprocessed using a customized version of the AFNI NHP preprocessing pipeline (Jung et al., 2021; Fujimoto et al., 2022). For each session, pre- and post-injection data were processed separately using the same parameters. In addition to a set of two dummy scans, the first three TRs of each EPI were removed to ensure that any magnetization effects were removed from the data before functional connectivity analyses. The images were first slice time corrected, then motion correction was applied, the EPIs were aligned to the within session T1-weighted image, and warped to the standard space. Following alignment to the standard space, the EPIs were blurred with an FWHM of 3 mm, and then converted to percent signal change. Finally, the motion derivatives from each scan along with cerebrospinal fluid and white matter signal were regressed out of the data. The residuals of this analysis were then used to compute the functional connectivity analysis described below.

#### Seed-based amygdala connectivity analyses

To examine the effects of drug on amygdala connectivity specifically, we created an amygdala mask from the standard D99 atlas (Saleem et al., 2021). We extracted the average time series from this amygdala ROI for each condition in each monkey, before and after drug treatment. We then calculated the Fisher’s z-transformed Pearson’s correlation between the average amygdala time series and the time series of every voxel in the brain (AFNI’s 3dTcorr1D). To statistically evaluate the effect of DCZ on amygdala functional connectivity, the correlation maps were submitted to an ANOVA with main effects of drug (VEH or DCZ), session, and monkey, with session nested under monkey.

#### Whole Brain MRI analyses

Whole brain connectome analyses were computed using 3dNetCorr function in AFNI (Cox, 1996; Taylor and Saad, 2013). For each animal and each session, we computed the correlation between ROIs of the D99 atlas (Saleem et al., 2021) and level three of the CHARM (Jung et al., 2021) and SARM (Hartig et al., 2021) atlases. The individual correlation matrices, or connectomes, were then Fisher’s z-transformed. For each session, we subtracted the pre-injection connectome from the post-injection connectome, creating a matrix of the functional connectivity changes. To statistically evaluate the effect of VEH or DCZ on the functional connectivity in across ROIs the difference connectomes were submitted to an ANOVA (function anovan in MATLAB) with main effects of drug and subject. ROI was modeled as a random effect, and session was nested under subject. We calculated the effect of CNO compared to VEH separately, as we only obtained one session CNO. Similar to the above analysis, the difference connectomes were submitted to an ANOVA with main effects of drug and subject, with ROI pair modeled as a random effect.

#### Seed to seed voxel based connectivity

We further examined the relationship between the amygdala and the ventrolateral prefrontal cortex on an individual voxel basis (Etzel et al., 2013). Using MATLAB and spm12, we extracted the individual time series from each voxel in the amygdala ROI and the vlPFC (ROI taken from the CHARM 6 atlas; combined masks for area 12o and area 12l; Jung et al., 2021). Analyses were performed separately for left and right hemispheres. A Fisher’s z-transformed correlation map was calculated between each voxel in the ipsilateral amygdala and the vlPFC. The average vlPFC z- value was then calculated for each amygdala voxel. The pre-injection correlation was subtracted from the post-injection matrix, producing a voxel-wise map of the change in functional connectivity between the two regions produced by drug administration.

### Neurophysiology data acquisition

We used the same anesthesia protocol as in the fMRI sessions, to maximize cross-modal comparisons. Following intubation, the head was fixed in a plastic stereotaxic frame and the sedation was maintained with low-level isoflurane throughout the session. In each session, the dura matter was penetrated using a tungsten guide tube (Crist Instruments, Hagerstown, MD) and then two (animal H) or three (animal L) 16-channel linear arrays (S-probe, Plexon, TX) were lowered using a NAN microdrive system (NAN instruments, Nazareth, Israel). In both animals, individual electrodes targeted both the basolateral amygdala and vlPFC in the same hemisphere (monkey H: left hemisphere, monkey L: right hemisphere), and an additional electrode (monkey L) targeted the contralateral amygdala. The recording depth and grid coordinates were calculated based on a high resolution T1w imaging with the animal in an MRI compatible stereotaxic frame. In animal H, the amygdala recording site was 22mm anterior to the interaural plane, 10mm to the right of the midline, and the vlPFC recording sites were 29.5±0.5mm anterior to the interaural plane, 14.5±0.5mm to the right of the midline. In animal L, the amygdala recording sites were 19mm anterior to the interaural plane, 7.5±0.5mm to the left of the midline and 19mm anterior to the interaural plane, 8.5±0.5mm to the right of the midline, and the vlPFC recording site was 27mm anterior to the interaural plane, 13.5±0.5mm to the left of the midline.

The electrodes were advanced to the target depths while listening to gray and white matter transitions to identify targets. We allowed neural signals to stabilize at the target depth for 1 hour before beginning data collection. Recording sessions consisted of three time periods: pre-injection (30 mins), injection (15-30 mins), and post-injection periods (30 mins). Following the pre-injection period, a drug (DCZ, CNO, or VEH) was injected intravenously, followed by a waiting period (injection) to allow for the known pharmacodynamics of the drugs in macaque monkeys (Nagai et al., 2020), and was consistent with the waiting period used during fMRI acquisition. The anesthetized recording was performed at least 1 week apart, and the order of drug conditions was counterbalanced across animals.

#### Single-unit data analysis

Wide-band signal was recorded for each electrode contact using an Omniplex system (Plexon, Dallas, TX) and stored for offline analysis. Signal was first band-passed (600 to 6000Hz) before extracting all spike waveforms exceeding 3 sd. Spikes from putative single neurons were then automatically clustered offline using the MountainSort plugin for MountainLab (Chung et al., 2017) and later curated manually based on the principal component analysis, interspike interval distributions, visually differentiated waveforms and objective cluster measures (Isolation probability > 0.75, Noise overlap probability < 0.2, Peak signal to noise ratio > 0.5 sd, Firing Rate > 0.05 Hz). Our dataset contained 155 amygdala (monkey H: 47; monkey L: 108) and 60 vlPFC neurons (monkey H: 39; monkey L: 21) across the 3 drug conditions. **Table 1** summarizes all recorded neurons across treatment conditions.

The spiking data was then analyzed using custom written MATLAB scripts (MathWorks, Natick, MA). Instantaneous firing rate for each neuron was extracted and binned into 30 second windows. Two periods were considered and defined as follows: the pre-injection period was from 10 min before and up until the injection timestamp, while the post-injection period started after the injection waiting period (15/30 minutes for DCZ/CNO) and ended 20 minutes later. The firing rate for each neuron was normalized using the mean and standard deviation during the pre-injection period (z-score normalization).

At the level of the neuronal populations, changes in firing rate were averaged for both recording period (pre-injection and post-injection) to calculate an average value for each neuron across each period. The difference between these averages (post-injection – pre-injection) was then statistically compared using Wilcoxon signed-rank test. Comparison between drug condition (VEH vs DCZ or VEH vs CNO) was performed on this same difference in averages using Kruskal-Wallis tests.

At the level of single neurons, we determined the proportion of neurons modulating their firing rate following drug injection. Neurons were considered modulated if their absolute normalized firing rate during the post-injection period was greater than 1.96sd for at least 4 time bins (equivalent of 2min out of the 20min post-injection period, 10%). To compare the proportion of modulated neurons across drug condition and area, we used mixed-effect logistic regressions, which included area (amygdala, vlPFC) and drug condition (VEH, DCZ or VEH, CNO) as factors, their interaction, and a random effect of animal (2 levels).

#### Local Field Potential analysis

The LFP data were analyzed using the FieldTrip toolbox (Oostenveld et al., 2011) and custom-made scripts on MATLAB. First, the wide band signal was subsampled at 1kHz before applying bipolar-referencing by subtracting the signal of every contact along each electrode array (16 contacts) with its closest neighbor (i.e., 15 bipolar sites). Visual inspection was used to reject bipolar sites containing noise from further analyses (across session average [min-max], 10% [0-20%] of bipolar sites).

To extract robust estimates of power modulation and functional connectivity, we first created 50 non-overlapping bins of 4 seconds each for both the pre-injection period and post-injection period, as defined above. This trial creation procedure and the following computations were repeated 100 times.

We applied a band-pass filter between 0.5 to 200Hz and an additional band-stop for line noise at 60Hz and its harmonics. We then performed a multitaper frequency transformation (7 tapers), with a spectral resolution of 0.5Hz (from 1 to 100Hz) and a spectral smoothing of +/-1Hz. This was used to estimate the power spectrum across areas, time periods and drug conditions. Grand-average power spectrograms in the post-injection period (across the 100 repetitions) were normalized using the average and standard deviation across the 100 pre-injection repetitions at every frequency bin (z-score normalization). Changes in normalized power in the post-injection periods were then assessed at every frequency bin using Wilcoxon signed-rank tests, while drug conditions effects (VEH vs DCZ or VEH vs CNO) were tested using Kruskal-Wallis tests. In both cases, we applied a threshold for significance at p=0.01 for 5 consecutive bins. Note that only amygdala recordings ipsilateral to the vlPFC recording sites in monkey L are reported, allowing for a direct comparison with the following coherency analyses.

The multitaper frequency decomposition output was also used to derive the imaginary part of the coherency between every pair of sites across the 50 trials for both time periods. Coherency values at each frequency bin were averaged across the 100 repetitions of the trial creation procedure, resulting in two tensors per recording session (one per time period) of size Nch x Nch x Nfq, where Nch is the number of channels and Nfq the number of frequency bins. For each of the 100 repetitions, we also created a matched null permutation where we scrambled the association between the different bipolar sites by randomizing the trial assignment for each site. This allowed us to create null coherograms, used to define our frequency band of interest. We further focused our analyzed on inter-areal coherency, specifically between ipsilateral amygdala-vlPFC pairs, therefore disregarding the contralateral amygdala in monkey L.

The existence of coherency at each frequency bin was established by comparing it to the null coherogram derived from permutations using Wilcoxon signed-rank tests (with a threshold for significance at p=0.01 for 5 consecutive frequency bins; **Supplemental Figure 3**). A frequency band of interest was further defined within the frequency bins showing significant coherencies by extracting the peak frequencies and the full width at a third of the peak maximum across both periods and all drug conditions for each monkey (monkey H: 6.5-14.5 Hz with peak at 10.5Hz, monkey L: 8-14Hz with peak at 9Hz). Given the overlap between monkeys, the frequency band used was based on the min and max frequencies across both monkeys (6.5 – 14.5 Hz), encompassing what is usually referred to as alpha oscillations.

Finally, we averaged the coherency observed between a given amygdala site and all possible vlPFC sites, resulting in two vectors of 6.5-14.5Hz coherency for each recording session, one for each period considered. To test for statistical difference between conditions and monkeys, we used two 2-way ANOVAs which included drug condition (VEH, DCZ or VEH, CNO) and monkey (H, L) as factors, as well as their interaction.

### Histological processing

Animals were deeply anesthetized with a sodium pentabarbitol solution and perfused transcardially with 0.1 M phosphate buffered saline (PBS) and 1% paraformaldehyde (PFA) solution followed by 0.1 M PBS and 4% PFA solution. Brains were removed and immersed in 4% PFA solution overnight. Following post-fixation, brains were transferred to a solution of 2% DMSO and 10% glycerol in 0.1 M PBS for 24-48 hours, then transferred to a solution of 2% DMSO and 20% glycerol in 0.1 M PBS for at least 24 hours. Cryoprotected brains were blocked in the coronal plane, then flash frozen in −80C isopentane. After flash freezing, the brains were removed, dried, and stored at −80C until sectioning. Tissue was cut serially in 50 um coronal sections on a sliding microtome (Leica SM 2010R) equipped with a freezing stage.

One series from each brain was mounted on glass slides, Nissl stained using cresyl violet solution, and coverslipped. A second series from each brain was used for confirmation of DREADD expression. Sections through the amygdala were selected and washed with PBS solution containing 0.3% Triton X-100 (TX-100). Endogenous peroxidase was quenched by incubation in 0.6% hydrogen peroxide in PBS for ten minutes. The sections were washed, then blocked for 1 hour in a solution of 1% bovine serum albumin, 1% normal goat serum, and 0.3% TX-100 in PBS. Primary antibody against the hemagglutinin tag (raised in rabbit, clone C29F4, 1:400, Cell Signaling Technology, Danvers, MA; cat # 3724, RRID:AB_1549585) was added to the blocking solution and sections were incubated overnight. At least one amygdala section for each animal, taken from a third series, was processed in a buffer solution overnight without primary antibody as a control. Incubation in primary antibody was followed by washes and a 2-hour incubation in anti-rabbit biotinylated secondary antibodies (1:200, Vector Laboratories, Burlingame, CA, RRID:AB_2313606). Sections were then washed and incubated for 1.5 hours in avidin-biotin complex solution (1:200; Vectastain standard kit, Vector Laboratories, RRID:AB_2336819). Finally, sections were washed, incubated in a 3-3’-diaminobenzidine tetrahydrochloride (DAB) solution for 10 minutes, and 0.006% hydrogen peroxide was added to the solution. Sections were carefully observed for a reaction and were placed in buffer solution after approximately 2.5-3.5 minutes, when staining was visually apparent. Sections were mounted on glass slides, air dried, and coverslipped with DPX mounting medium.

Images of sections stained for DREADD receptors were acquired using a Zeiss Apotome 2 microscope equipped with a Q-Imaging digital camera, motorized stage, and Stereo Investigator software (MBF Bioscience). Images were acquired using consistent exposure time and brightness settings within the same section. Any adjustments made afterward (e.g. to increase brightness) were applied uniformly to the image. To obtain wide field images of the entire section, the outline of the tissue was defined using a 5x lens. Tiled images of a single section were obtained sequentially, constrained by the boundaries of the tissue, and a 10% overlap and image stitching was used to create a composite image of all tiled photos.

## Data Availability Statement

The data that support the findings of this study are available from the corresponding author upon reasonable request.

## Acknowledgements

[AF, CE, FMS, NB, BER and PHR are supported by grants from NIMH and the BRAIN initiative (R01MH110822 and RF1MH117040). BER is supported by grants from NIMH (R01MH111439). AF is supported by Overseas Research Fellowship from Takeda Science Foundation and a Brain & Behavior Research Foundation Young Investigator grant (#28979). SHF is supported by a grant from the Hope for Depression Research Foundation. We would like to thank Dr Paula Croxson for providing the foundation on which this work was built and Jairo Munoz for assistance with data acquisition. We thank Dr Paul Taylor for help with fMRI data pre-processing. For help with fMRI analysis we thank Drs Alex Franco and Vincent Costa. We also thank Dr Jian Jin for providing clozapine N-oxide powder and Drs Naohisa Miyakawa and Vincent Costa for their advice on setting up S-probe configuration. We thank Dr David Leopold for his advice and suggestions on the manuscript. Finally, we thank the veterinary and animal care staff at Mount Sinai for their expertise and support.]

## Notes

**Conflict of interest** The authors declare no competing interests.

### Competing Interest Statement

The authors have declared no competing interest.

### Summary of Updates

We have corrected NIH grant information in the acknowledgements and changed the creative commons license in this revision.

