## Supplemental Figures for "The neural basis of resting-state fMRI functional connectivity in fronto-limbic circuits revealed by chemogenetic manipulation"

+ Joint last author

**This file includes:**

Supplementary Figures 1-3

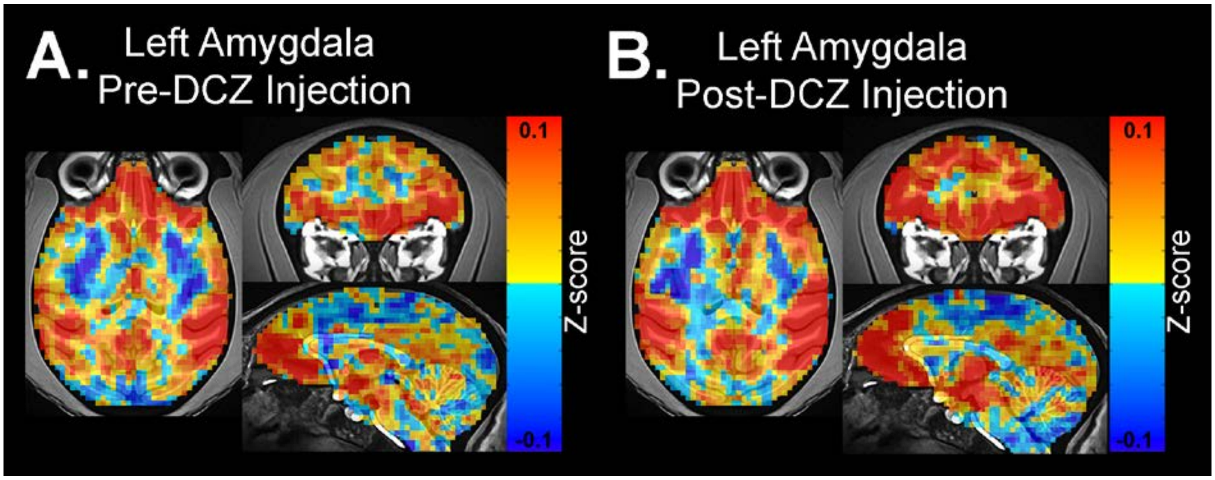


**Supplemental Figure 1. Representative changes in functional connectivity between amygdala and frontal cortex, unthresholded connectivity maps.** Changes in FC with a left amygdala seed region in animal L in a single session, before and after DREADD activation with DCZ. Pre-DCZ injection period **(A)** and post-injection period **(B)**. Scale bar indicates z-score of rs-FC. No threshold or clustering has been applied.

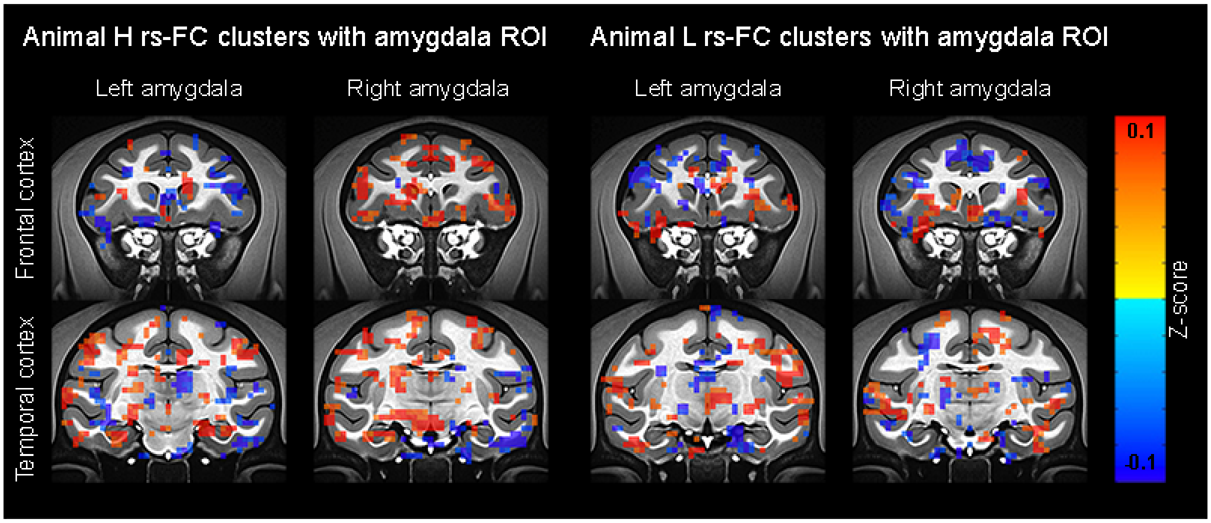


**Supplemental Figure 2. Change in amygdala ROI rs-FC with frontal and temporal cortex after amygdala inhibition.** Example coronal sections from each animal showing the change in rs-FC with a left or right amygdala seed after DREADD activation with DCZ, as compared to VEH. Threshold p=0.05, cluster size ≥5 voxels, voxel faces touching.

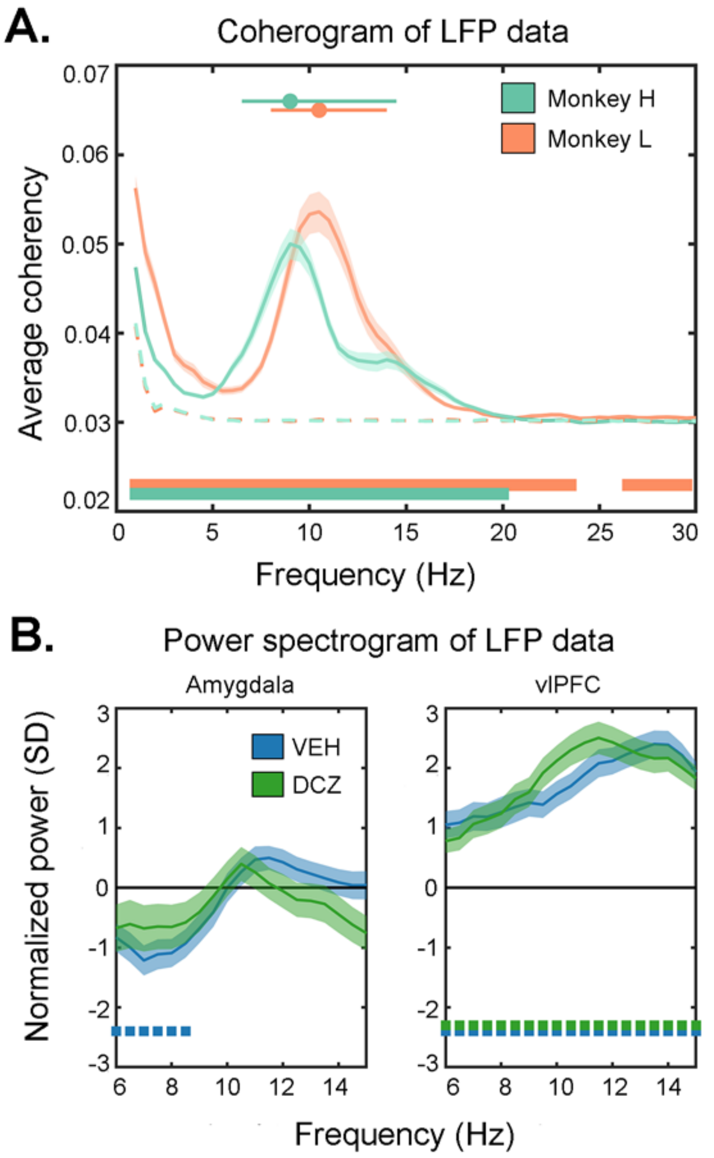


**Supplemental Figure 3. Across the alpha band, coherency is increased, but power is unaltered. A**) Average coherency across all periods and drug conditions for both animals. Bottom lines indicate difference in coherency from randomized permutations. Lines/circles (top) show the peak frequency + 1/3$^{rd}$ of the max value. **B**) Post-injection power normalized to the pre-injection period, across the frequency band of interest defined in A, for the amygdala (left) and vlPFC (right) after treatment with DCZ or VEH. Dotted line (bottom) shows significant differences from pre-injection power (Wilcoxon signed rank test, $p<0.01$ for 5 consecutive bins). No significant differences between DCZ and VEH were observed in either region (Kruskall-Wallis test, $p<0.01$ for 5 consecutive bins).
